# Diazepam attenuates the effects of cocaine on locomotion, 50-kHz ultrasonic vocalizations and phasic dopamine release in the nucleus accumbens of rats

**DOI:** 10.1101/2020.10.13.338343

**Authors:** William N. Sanchez, Jose A. Pochapski, Leticia F. Jessen, Marek Ellenberger, Rainer K. Schwarting, Donita L. Robinson, Roberto Andreatini, Claudio Da Cunha

**Author notes:** Correspondence Claudio Da Cunha, Laboratório de Fisiologia e Farmacologia do Sistema Nervoso Central, Department of Pharmacology, Universidade Federal do Paraná, Curitiba, PR, Brazil. Funding information Coordination for the Improvement of Higher Education Personnel – Brazil (CAPES) – Finance Code protocol 88881.198683/2018-01 and by The Brazilian National Council for Scientific and Technological Development (CNPq) grant process number 432061/2018-5.

## Abstract

**Background and Purpose:** Currently, no effective drug exists to treat cocaine use disorders, which affect millions of people worldwide. Benzodiazepines are potential therapeutic candidates, as microdialysis and voltammetry studies have shown that they can decrease dopamine release in the nucleus accumbens of rodents. In addition, we have recently shown that diazepam blocks the increase in dopamine release and the affective marker 50-kHz ultrasonic vocalizations (USV) induced by DL-amphetamine in rats.

**Experimental Approach:** Here we tested whether administration of 2.5 mg·kg^−1^ diazepam (i.p.) in adult male Wistar rats could block the effects of 20 mg·kg^−1^ cocaine (i.p.) on electrically evoked phasic dopamine release in the nucleus accumbens measured by fast-scan cyclic voltammetry, as well as 50-kHz USV and locomotor activity.

**Key Results:** Cocaine injection increased evoked dopamine release up to 3-fold within 5 min and the increase was significantly higher than baseline for at least 90 min. The injection of diazepam 15 min later attenuated the cocaine effect by nearly 50% and this attenuation was maintained for at least 30 min. Stimulant drugs, natural rewards and reward predictive cues are known to evoke 50-kHz USV in adult rats. In the present study, cocaine increased the number of 50-kHz USV of the flat, step, trill, and mixed kinds by 12-fold. This effect was at maximum 5 min after cocaine injection, decreased with time and lasted at least 40 min. Diazepam significantly blocked this effect for the entire duration of the session. The distance travelled by control rats during a 40-min session of exploration in an open field was at maximum in the first 5 min and decayed progressively until the end of the session. Cocaine-treated rats travelled significantly longer distances when compared to the control group, while diazepam significantly attenuated cocaine-induced locomotion by up to 50%.

**Conclusions and implication:** These results suggest that the neurochemical, affective, and stimulant effects of cocaine can be mitigated by diazepam.

**What is already known:**

- Diazepam decreases dopamine release in the rodent nucleus accumbens (NAc) and reduces some effects produced by DL-amphetamine.

**What this study adds:**

- Diazepam attenuated the increase in phasic dopamine release caused by cocaine.
- Diazepam blocked the effect of cocaine on 50-kHz USV and locomotor activity.

**Clinical significance:**

- This study demonstrates that diazepam can block specific effects of cocaine that likely contribute to addiction.

## 1 INTRODUCTION

Although cocaine use disorders are a global epidemic problem that is growing in many parts of the world, there is no FDA approved pharmacological treatment for this disorder (Farrell et al., 2019; Kampman, 2019, UNODC 2020). Thus, it is urgent to find an effective and safe pharmacotherapy to treat cocaine and amphetamine use disorders.

Cocaine binds to the dopamine transporter, limiting protein movement and acting as a dopamine transporter blocker (Wang et al., 2015). Due to this interaction, cocaine causes an increase in extracellular dopamine levels in the dorsal and ventral striatum (Church et al., 1987; Hernandez and Hoebel, 1988; Garris and Wightman, 1995; Heien et al., 2005). The increase in the intensity and frequency of dopamine release in the nucleus accumbens (NAc) caused by cocaine reinforces self-administration of this drug and increases locomotor activity in rodents and humans (Tilley et al., 2007; Hall et al., 2009). 50-kHz USV emitted by adult rats in rewarding situations or in response to cues with incentive salience are also evoked by cocaine and other stimulants in adult rats (Simola et al., 2012a, 2014). In addition, sensory cues paired with cocaine acquires incentive salience and later can elicit dopamine release causing drug-seeking behaviors (Phillips et al., 2003). Therefore, it is possible that blocking the cocaine amplification of dopamine release can be a strategy to block the rewarding and motivational effects of this drug (German et al., 2015; Volkow et al., 2017).

Diazepam is a benzodiazepine used to treat anxiety, alcohol withdrawal, skeletal muscle spasm and convulsive disorders (Calcaterra and Barrow, 2014). Diazepam binds to an allosteric site in the GABA_A_ receptor (GABA_A_-R), augmenting the activity of the gamma-aminobutyric acid (GABA) transmission (Zhu et al., 2018; Masiulis et al., 2019) by enhancing the GABA-A receptor affinity for GABA and subsequently increasing the channel-open frequency (Uusi-oukari and Korpi, 2010). Diazepam has been shown to promote cocaine abstinence (Augier et al., 2012) and to attenuate the stimulant locomotor effect of morphine and amphetamine (Panhelainen et al., 2011). Also, we previously reported that diazepam blocks the increase in phasic dopamine release in the mouse NAc and rat 50-kHz USV, both evoked by amphetamine (Gomez-A et al., 2017; Guaita et al., 2018). As both amphetamine and cocaine increased dopamine release, it is likely that diazepam will similarly affect cocaine-induced dopamine release. However, as mentioned above, the mechanisms by which amphetamines and cocaine increase dopamine release are not identical. Therefore, in the present study we tested the hypothesis that diazepam blocks the effect of cocaine on dopamine release. Here we used in vivo fast scan cyclic voltammetry (FSCV) to show that diazepam can mitigate the increase in the electrically evoked phasic dopamine release in the rat NAc. We also show that diazepam attenuated the increase in the number of rat 50-kHz USV and the increase in locomotion observed in rats treated with cocaine.

## 2 METHODS

### 2.1 Animals

Adult male Wistar rats (n=31 for FSCV, n=41 for USV) were housed in groups of five in polypropylene cages (41 cm × 34 cm × 16 cm) with sawdust bedding under a 12 h/12 h light/dark cycle (lights on at 7:00 a.m.) and a controlled temperature (22 ± 2 °C). Food and water were available ad libitum. Males were used to better compare to previous studies (Gomez-A et al., 2017, Guaita et al., 2018). All procedures were in accordance with the National Institutes of Health Guide for the Care and Use of Laboratory Animals and approved by the institutional Ethics Committee for Animal Experimentation of the Universidade Federal do Paraná (Protocol 1157) consistent with Brazilian law (Bil#11.794/8 October 2008).

### 2.2 Chemicals and solutions

All chemicals were purchased from Sigma Aldrich (St Louis, MO), except for cocaine HCl (donation from Curitiba Police Department inquiry # 5016610-91.2019.4.04.7000/PR). Cocaine was 98% pure, as determined by nuclear magnetic resonance. Cocaine HCl was dissolved in saline (0.9 % w/v) to 20.0 mg·mL^−1^ and diazepam was dissolved in the vehicle solution (2% propylene glycol, 8 % ethyl alcohol, using 5% sodium benzoate and benzoic acid as buffers, and 2% benzyl alcohol, pH= 7.4) to 5 mg·mL^−1^.

### 2.3 *In vivo* Fast Scan Cyclic Voltammetry

Animals were anesthetized with urethane (1.5 mg·kg^−1^ i.p.) and placed in a stereotaxic frame (David Kopf Instruments, Tujunga, CA). Urethane was used because there is no effect on dopamine clearance *in vivo* (Garris et al., 2002; Sabeti et al., 2003). A stimulation electrode (Plastics One, Roanoke, VA, USA, polished tips, 1 mm apart) was implanted in the ventral tegmental area (VTA) (AP= −0.52, ML= + 0.10, DV= −0.82) and a carbon-fiber electrode (10-80 μm length, 7 μm diameter, Goodfellow, Huntingdon, England) was implanted in the NAc (AP= + 0.19, ML= + 0.14, DV= −0.64) according to a rat brain atlas (Paxinos and Watson, 1998). An Ag/AgCl reference electrode was placed in the contralateral hemisphere and affixed with dental cement.

The parameters of electrical stimulations and neurochemical recordings were as previously described (Gomez-A et al., 2017) with some modifications. In brief, every 100 ms, a triangular waveform potential (−0.4 V to +1.3 V to −0.4 V vs Ag/AgCl) was applied at a rate of 400 V/s. After 30 min of waveform application to condition and stabilize the electrode, trains of 35 biphasic pulses (0.5 ms per pulse, 600 μA, 60 Hz, 24 pulses) were applied to the stimulating electrode every 300 s with a programmable optical isolator pulse generator (MINCS, Mayo Investigational Neuromodulation Control System, Mayo Clinic). FSCV measurements were recorded with a Wireless Instantaneous Neurotransmitter Concentration Sensor system (WINCS, Mayo Clinic) and processed with the WINCS and MINCS software (version 2.10.4.0, Mayo Clinic). Five trains of electrical pulses were applied to confirm the stability of the FSCV recording. Then, another six trains of pulses were applied (300 s apart) and the evoked currents were taken as baseline, given that the evoked dopamine release did not vary by more than 16%. Immediately thereafter, rats received cocaine (20.0 mg·kg^−1^) or saline and another five trains of pulses were applied. Next, diazepam (2.5 mg·kg^−1^) or vehicle was administered and electrochemical data acquisition continued for 70 min with electrical stimulation every 300 s.

The background-subtracted cyclic voltammograms were obtained by subtracting voltammograms collected during stimulation from those collected up to 2 s before the stimulation. Voltammetric responses were displayed as pseudo-color plots with the abscissa as the voltage, the ordinate as the acquisition time, and the current encoded in color. Temporal responses were determined by monitoring the current at the peak oxidation potential for dopamine in successive voltammograms. To estimate dopamine concentration, the recording electrodes were calibrated using a flow-cell (dopamine solution of 20 μM in TRIS buffer) and the calibration factor was 1nA = 1.033 ± 0.407 μM. To confirm the identity of the evoked dopamine release, background-subtracted cyclic voltammograms of the *in vivo* electrochemical signal immediately after stimulation were averaged and correlated with voltammograms from the *in vitro* calibration (R^2^=0.963, p <0.001).

### 2.4 Histology

After FSCV recording, rats were decapitated and brains were fixed in 4% formaldehyde for 10 days. Coronal slices of 50 μm were obtained using a vibratome (VT 1000S, Leica Biosystems), stained with thionine, and compared to the rat atlas of Paxinos and Watson (1998) to locate the electrodes’ track.

### 2.5 Ultrasonic vocalizations and locomotor activity

Animals were divided in four groups that received: (i) saline and the vehicle; (ii) saline and diazepam (2.5 mg·kg^−1^ dissolved in vehicle); (iii) vehicle and cocaine (20.0 mg·kg^−1^ dissolved in saline); (iv) cocaine (20.0 mg·kg^−1^) and diazepam (2.5 mg·kg^−1^). Rats were tested in two sessions: baseline and treatment, 40 min each, 24 h apart. Drugs were administered only immediately before the treatment session. In each session, each animal was gently placed into an acrylic box (40 × 40 × 40 cm) with fresh sawdust bedding on the floor. USV were recorded using a microphone (UltraSoundGate Condenser Microphone, CM16; Avisoft Bioacoustics, Berlin, Germany) placed in the center and 25 cm above the floor of the box, controlled by the Avisoft Recorder 2.7 software. USV 50-kHz calls were counted and classified using DeepSqueak 2.6.1 software (Coffey et al., 2019). The videos recorded during USV sessions were used to measure locomotor activity using automatized analysis with EthoVision® XT software (Noldus, Wageningen, Netherlands).

### 2.6 Statistics

We applied the D’Agostino & Pearson and Shapiro-Wilk tests of normal distribution. Variability among rats due to differences in electrode location and length was reduced by using the equation “Y= [I_i_/I_BL_] * 100”, where I_i_ is the oxidation current on time i, and I_BL_ is the baseline evoked dopamine release. In each rat, the I_BL_ was calculated by averaging the heights of the first 6 peaks of I_i_ recorded after the oxidation current values had stabilized (see 2.3 for details). As neither raw data nor the transformed data passed the tests of normal distribution, we used a generalized linear regression (GLM) model with Poisson-distribution, link log function and Wald chi-square comparisons to test time and drug effects. When appropriate, Sidak post hoc tests were used to compare differences among groups. We used odds ratios and confidence intervals to describe the magnitude of differences between groups and times.

The total number of USV calls (number of calls) did not pass the D’Agostino & Pearson and Shapiro-Wilk tests of normal distribution. However, after transformation with the algorithm [Y = sqrt(Y)], they passed these tests of normal distribution and were analyzed by two-way ANOVA with multiple comparisons and repeated measures, followed by the Tukey test to measure differences between baseline and treatment for each group.

Locomotion scores over time passed the normal distribution tests and were analyzed by repeated measure ANOVA followed by the Tukey test. The Poisson GLM was performed by using SPSS statistic software (IBM SPSS Statistics for Windows, Version 26.0. Armonk, NY: IBM Corp.). All the other analyses were done using GraphPad Prism software (San Diego, CA). The 5 min-bin graphs of FSCV were made using Python (Python Software Foundation. Python Language Reference, version 3.7, Amsterdam, Netherlands) and the other graphs were made using GraphPad Prism software. Differences among groups were considered significant if p < 0.05.

## 3 RESULTS

### 3.1 Diazepam attenuated the cocaine effect on the phasic release of dopamine in the NAc

To evaluate whether i.p. injection of diazepam blunted the effect of cocaine on phasic dopamine release, we compared evoked dopamine release in three groups of rats: cocaine+diazepam, cocaine+vehicle, and saline+vehicle (Figure 1a). The placements of the stimulating electrodes in the VTA and the recording electrodes in the NAc were confirmed (Figure 1b). Cocaine increased electrically evoked dopamine release, an effect that persisted for at least 75 min and was attenuated by diazepam (Figure 1c). The generalized linear model yielded significant main effects of group (Wald chi-square 1663.2, df=2, p <0.001) and time (Wald chi-square 987.6, df=3, p <0.001), as well as a significant interaction of group by time (Wald chi-square 765.6, df=6, p <0.001). Thus, we compared parameter estimates within groups across time points and between groups at each time point (see Figure 1c for post-hoc comparisons). Evoked dopamine in the saline+vehicle group did not differ across the experiment. In contrast, dopamine release in the cocaine+vehicle and cocaine+diazepam groups increased similarly after cocaine injection (0-25 time bin). While the cocaine+vehicle group continued to exhibit high levels of evoked dopamine release at each post-cocaine time bin, administration of diazepam in the cocaine+diazepam group significantly reduced this cocaine effect on dopamine release. Figure 1d shows representative examples of voltammograms and color plots taken from one rat. Note that after the electrical stimulation, current increased only in the potentials near the optimal dopamine oxidation potential. These examples also illustrate that, at the dopamine oxidation potential, the evoked current increase was higher after cocaine injection and returned back to a lower level after the diazepam injection.

**FIGURE 1.**
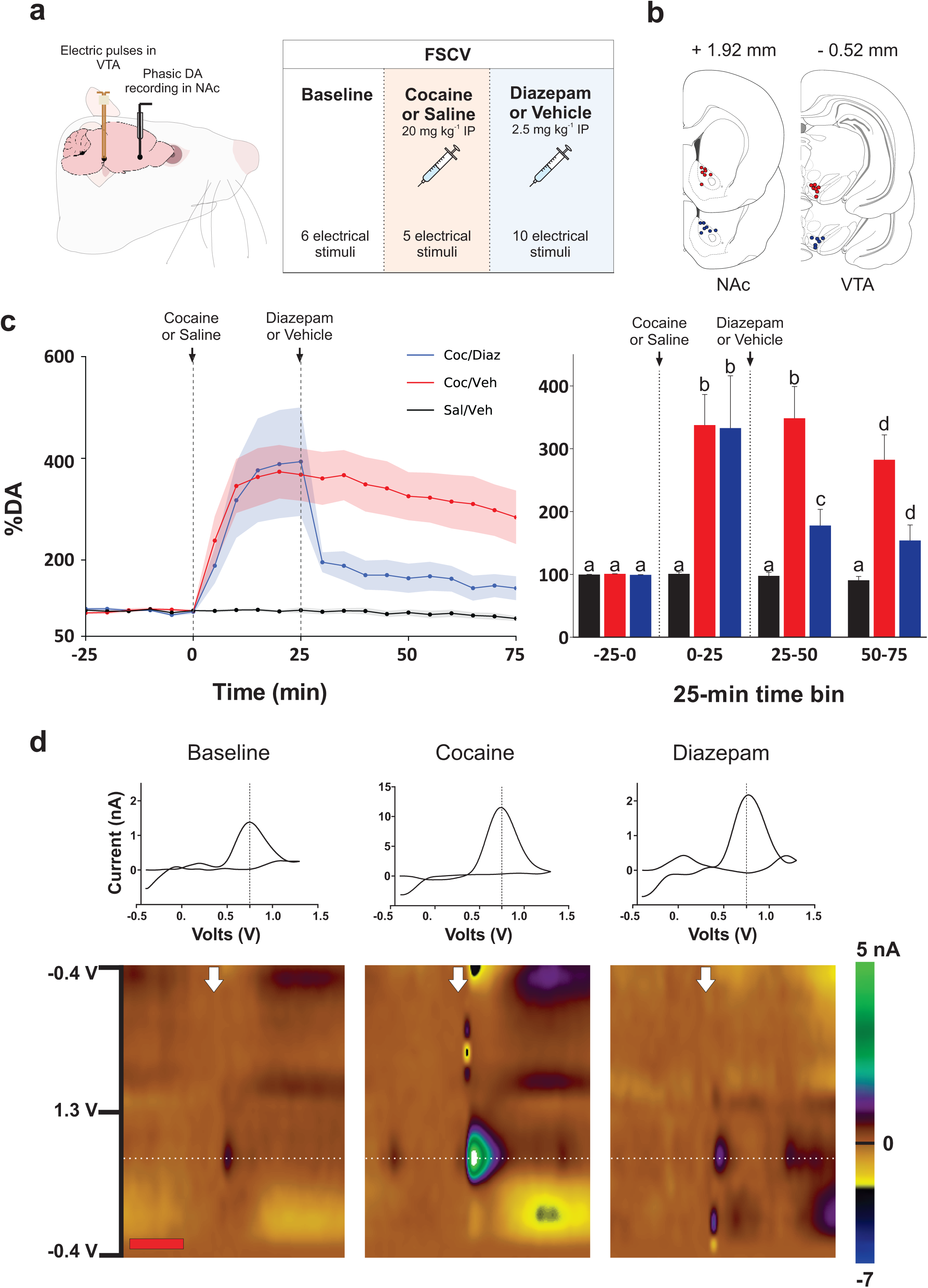
Diazepam blocks cocaine effect on dopamine release in the NAc evoked by electrical stimulation of the VTA. **a**. Experiment design, **b**. Schematic drawing for electrode implantation; red dots indicate electrode implantation for animals treated with cocaine and dark blue dots indicate electrode implantation in cocaine + diazepam group, **c**. Average results of diazepam effect on NAc Da release in rats treated with saline+vehicle (n=8), cocaine+vehicle (n= 7) and cocaine+diazepam (n= 7) recorded every 5 min for 105 min (left) and 25 min bins (right). Shading indicates ±SEM. Data were analyzed with GLM Poisson regression with post hoc Tukey tests. Means with different letters are significantly different from each other (p < 0.05), and means with the same letter are not significantly different. **d**. Representative examples taken from an individual rat of background-subtracted cyclic voltammograms and color plots of the diazepam effect. In voltammograms and color plots, the dotted line indicates maximum current potential (0.75 V); note the difference in current scale for the voltammograms. For color plots, the red bar indicates five seconds and the white arrow indicates the stimulation time.

### 3.2 Diazepam blocked the cocaine effect on 50 kHz USVs

It is well established that psychostimulants produce an increase in appetitive USV in rats, which can serve as index of positive affect (Simola et al., 2012a, 2014). Therefore, we evaluated if diazepam would reduce such an effect (Figure 2). To address this question, we placed adult male rats in an open field to record USV for 40 min on two days (Figure 2a). On the first day we recorded basal USV (baseline day) and on the second day we recorded USV under the effect of diazepam, cocaine, and saline+vehicle treatments at the same doses as used in the FSCV experiment. A repeated measures ANOVA showed a significant session factor (F_1,36_ = 23.08, p < 0.001), a significant group factor (F_3,36_ = 6.32, p < 0.01) and a significant interaction between these factors (F_3,36_ = 13.29, p < 0.001). No significant differences among groups were detected on the baseline day (data not shown). We used Tukey test in all post-hoc comparisons among groups. On the test day, only animals that received cocaine presented an increase in the 50-kHz USV compared with the other groups (p < 0.001) and compared to the baseline day (p < 0.001). Diazepam blocked this cocaine effect. No significant difference was observed between the group that received cocaine+diazepam and the group that received saline+vehicle (p = 0.99). In addition, no significant difference was observed between the group that received cocaine+diazepam and the group that received saline+diazepam (p=0.49, Figure 2b).

**FIGURE 2.**
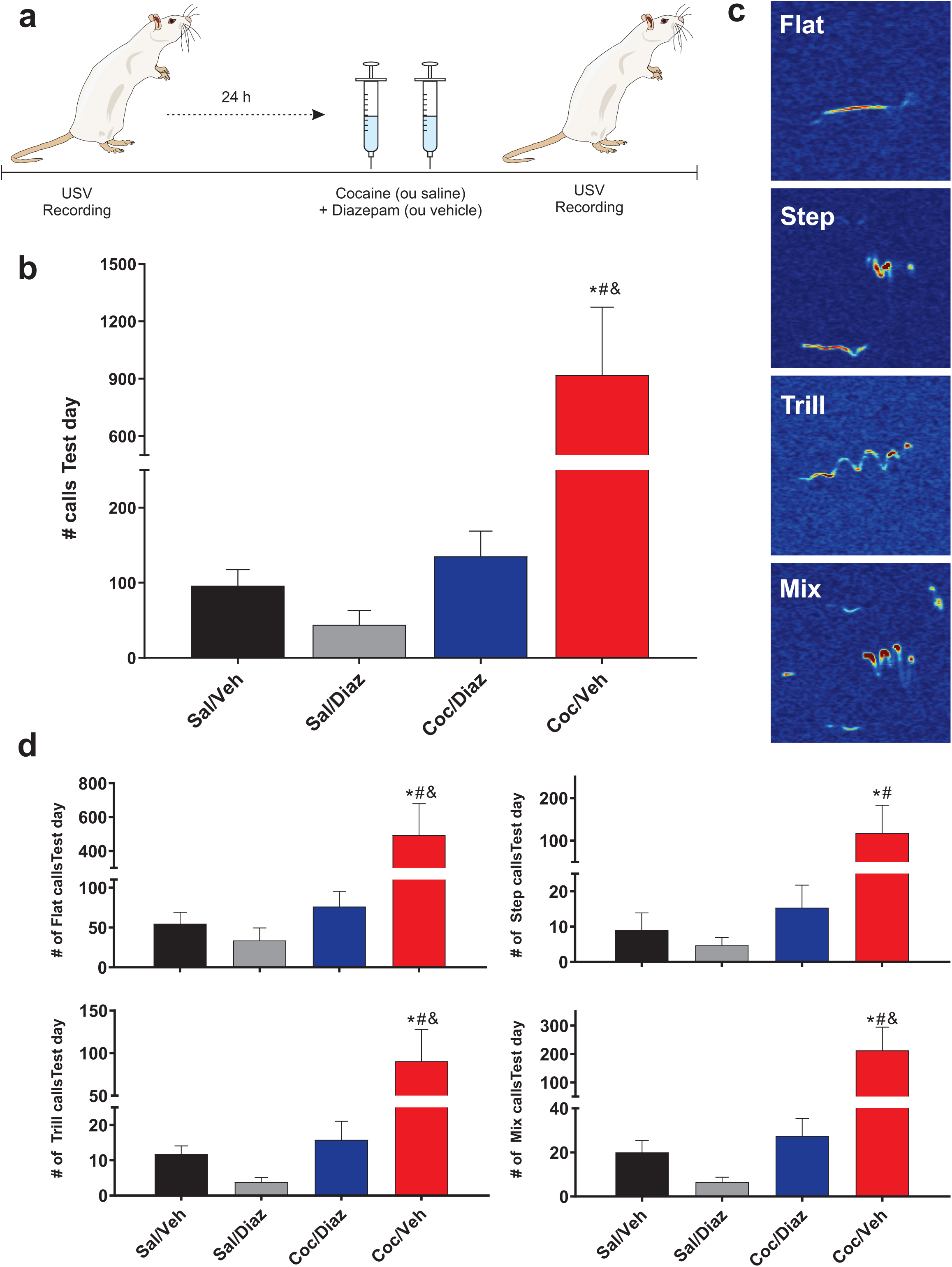
Diazepam blocked the cocaine-induced 50-kHz USV. **a**. Experimental design. **b**. Diazepam effect on total 50-kHz calls on the test. **c**. Examples of USV call type. **d**. Call subtypes on the test day (flat: upper left, step: upper right, trill: lower left, mix: lower right). * p < 0.05 compared to the Sal/Veh group, # p < 0.05, compared to the Sal/Diaz group, & p < 0.05 compared to the Coc/Diaz group, Tukey test after two-way ANOVA. Sal/Veh (n=10), Sal/Diaz (n=10), Coc/Veh (n=11), Coc/Diaz (n=9). Coc, 20 mg·kg^−1^ cocaine; Diaz, 2.5 mg·kg^−1^ diazepam, Sal, saline (cocaine’s vehicle), Veh, diazepam’s vehicle.

We also tested the effects of cocaine and diazepam on four subtypes of USV calls: flat, step, trill and mix (Figure 2c). Cocaine-treated rats presented higher numbers of all call subtypes (p < 0.01) compared with the saline+vehicle group. Diazepam blocked the cocaine effect on flat, trill and mix, subtypes: the number of calls emitted by the rats that received diazepam and cocaine was significantly different compared to the rats that received cocaine. In addition, the number of flat, trill and mix 50 kHz USV emitted by the cocaine+diazepam group was not significantly different compared to the saline+vehicle group (P = 0.98). The effect of diazepam on the step calls was less clear. The result suggests that diazepam attenuated the effect of cocaine because the number of step calls of the cocaine group, but not of the cocaine+diazepam group, was significantly different compared to the saline+vehicle group (p < 0.05). However, the number of step calls in the cocaine+diazepam group was not significantly different compared to the cocaine group (Figure 2d). Figure 3a shows that the saline+vehicle and saline+diazepam groups emitted 50 kHz USV mainly in the first 5 min and only few calls during the rest of the session. The animals of the cocaine+vehicle group presented a different distribution; they emitted calls throughout all the test session with a decay in call number across time. In contrast, animals treated with both diazepam and cocaine presented a similar pattern of call distribution as the saline+diazepam and saline+vehicle groups. One-way ANOVA showed that neither diazepam, cocaine, nor their combination had an effect on call duration (Figure 3b, F_3, 36_ = 0.495, p = 0.69) and call frequency (Figure 3c, F_3,,36_ = 0.88, p = 0.46).

**FIGURE 3.**
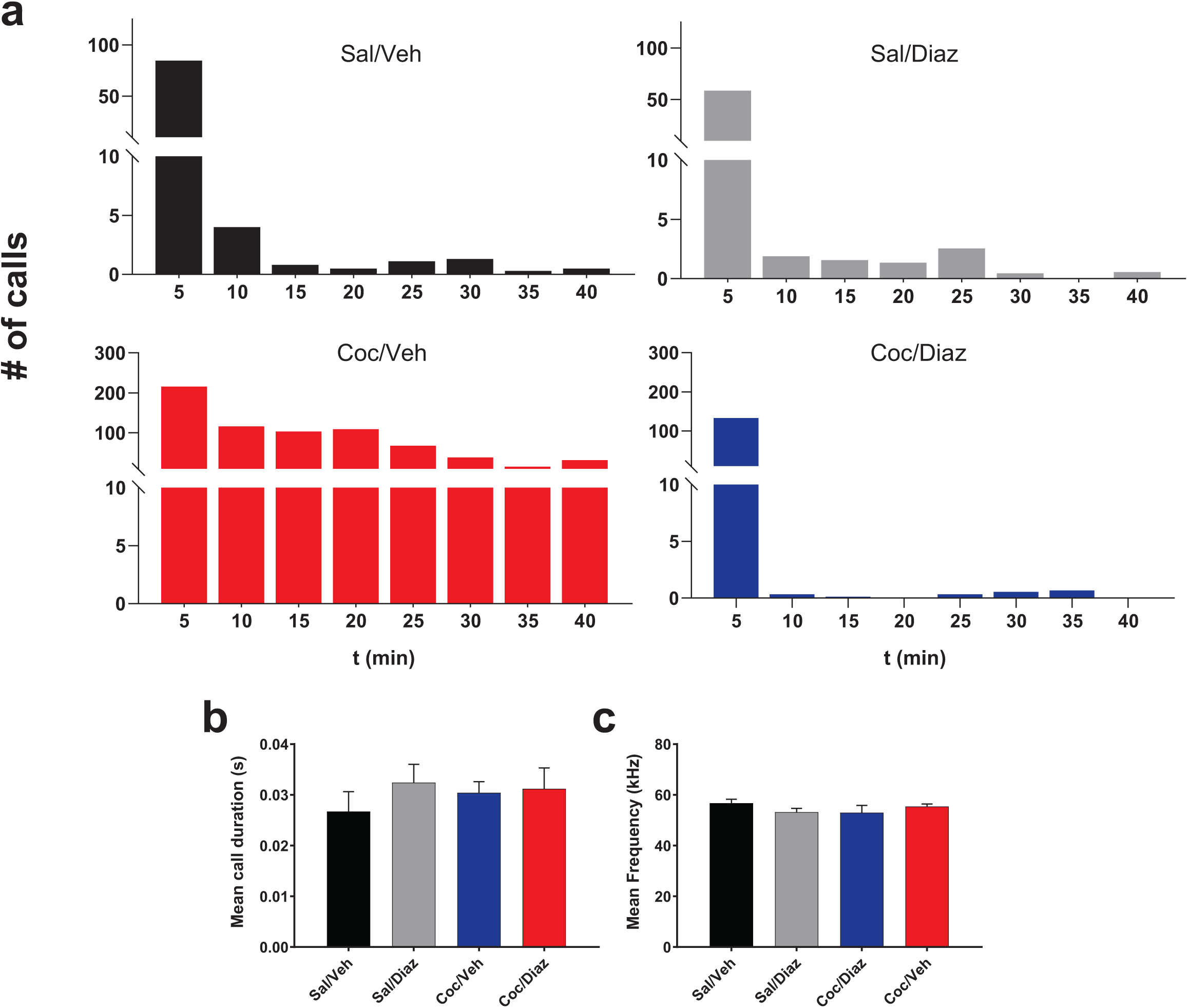
Time course and acoustic parameters of the cocaine and diazepam effects on appetitive 50-kHz USV. Sal/Veh (n=10), Sal/Diaz (n=10), Coc/Veh (n=11), Coc/Diaz (n=9). Coc, 20 mg·kg^−1^ cocaine; Diaz, 2.5 mg·kg^−1^ diazepam Sal, saline (cocaine’s vehicle), Veh, diazepam’s vehicle.

### 3.4 Diazepam decreases locomotor activity produced by cocaine

Locomotor activity (travelled distance) was measured in the same baseline and test sessions in which USV were recorded (Figure 4). A repeated measures ANOVA showed a significant session factor (F_1,16_= 30.18, p < 0.001), a significant group factor (F_3,36_ = 38.86, p < 0.001) and a significant session x group interaction (F _3,36_ = 40.3, p < 0.001). We used Tukey test in all post-hoc comparisons among groups. No significant difference among groups was observed on the baseline day (P > 0.99, data not shown). Only rats that received cocaine on the test day travelled longer distances compared to the same group on the baseline day and compared to all other groups on the test day (p < 0.05). As shown in Figure 4a, on the test day both the group treated with cocaine and the group treated with cocaine+diazepam travelled longer distances compared to the saline+vehicle group (p < 0.05). However, the distance traveled by the cocaine+diazepam group was significantly shorten compared to the cocaine group (p <0.05). This finding suggests that diazepam blocked partly, but not completely, the stimulant cocaine effect on locomotion. Representative path tracks are shown in Figure 4b. Figure 4c shows the average distance traveled by rats during the 40 min of the test sessions divided into 5-minute bins. Note that while the locomotor activity of the control group decreased 5 min after the start of the test session, the locomotor activity of the rats treated with cocaine increased at that time and this increase was maintained for the rest of the session. Note also that the co-administration of diazepam attenuated this locomotor stimulant effect of cocaine during the 40 min of the test session.

**FIGURE 4.**
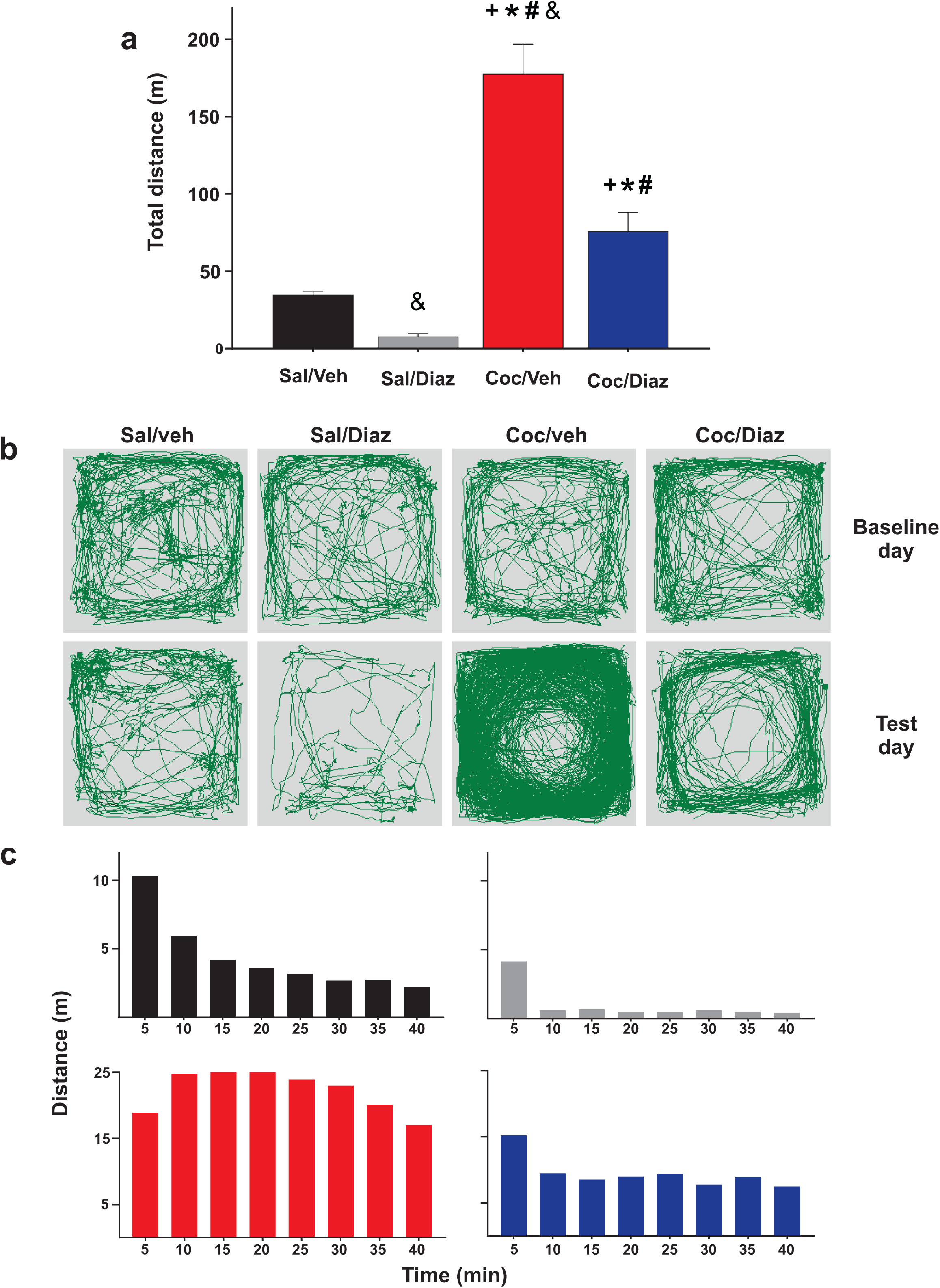
Diazepam attenuates the cocaine effect on locomotion activity. **a**. Travelled distance during the 40 min session. **b**. Representative individual pathways of locomotion for baseline and test day. **c**. Travelled distance in 5-min intervals. + p < 0.05 compared to the same group on the baseline day; * p < 0.05, compared to saline vehicle in the test day; # p < 0.05, compared to saline+diazepam on the test day; & p < 0.05 compared to cocaine+diazepam on the test day, Tukey test after ANOVA. Coc, 20 mg·kg^−1^ cocaine; Diaz, 2.5 mg·kg^−1^ diazepam; Sal, cocaine vehicle (saline); Veh, diazepam vehicle.

## 4 DISCUSSION

To evaluate the potential of benzodiazepines to treat stimulant drug disorders, it is important to determine whether benzodiazepines can counteract the neurochemical and behavioral effects of a stimulant drug. It have previously been shown that diazepam can attenuate the effects of amphetamine on NAc dopamine release, measured by FSCV and microdialysis (Zetterström and Fillenz, 1990; Murai et al., 1994; Gomez-A et al., 2017). In another study, we showed that diazepam can also attenuate the effects of amphetamine on rat 50 kHz USV and stereotyped behavior (Guaita et al., 2018). In the present study, we used FSCV to show that systemic injection of diazepam not only decreases dopamine release, but also reverts the increase in dopamine release in the NAc caused by cocaine in rats. Additionally, we showed that diazepam blocked the increase in the emissions of 50 kHz USV calls and attenuated the increase in locomotor activity caused by cocaine in rats. Together, these data indicate that benzodiazepines can blunt multiple key effects of acute stimulants.

In the present study we also tested the effect of diazepam on cocaine-evoked 50 KHz USV, a rat model of the rewarding effect of stimulant drugs. Rats emit 50 KHz USV in response to rewarding stimuli (Burgdorf et al., 2008; Brenes and Schwarting, 2015; Brenes et al., 2016; Wendler et al., 2016a; Engelhardt et al., 2017; Simola and Brudzynski, 2018) and in response to reward-predictive stimuli (Brenes and Schwarting, 2015). Dopamine is released in the NAc in response to the same stimuli (Ikemoto et al., 2015; Vega-Villar et al., 2019). Likewise, rats vocalize at 50-kHz USV when challenged with cocaine (Barker et al., 2010; Browning et al., 2011), amphetamine (Burgdorf et al., 2001; Pereira et al., 2014; Wöhr et al., 2015; Guaita et al., 2018), and other stimulant drugs (Simola et al., 2012b; Wendler et al., 2016b; Simola and Brudzynski, 2018). Furthermore, rat 50 kHz USV can be elicited by dopamine receptor agonists (Brudzynski et al., 2012), and stimulation of the VTA (Scardochio et al., 2015). Conversely, 50-kHz USV are reduced by VTA lesions and by dopamine receptor antagonists (Burgdorf et al., 2007; Ringel et al., 2013; Brudzynski, 2015; Rippberger et al., 2015).

When an individual self-administers stimulant drugs, the self-administration behavior that preceded the drug-induced dopamine release is reinforced. This self-sustained cycle is thought to play a key role in drug addiction. Furthermore, because these drugs cause dopamine release, the stimuli that were paired with the drug acquire incentive salience.

Over time, stimuli that became predictive of drug effects acquire the property to cause dopamine release in the striatum, which increases the motivation (wanting) that acts as an incentive for drug-seeking and drug consumption (Volkow et al., 2017). In agreement with this hypothesis, it has been shown that rats responded to a cue that precedes access to cocaine by exhibiting 50-kHz USV and that the number of calls were exacerbated when rats were abstinent (Maier et al., 2010). Therefore, the finding that diazepam blocks the effects of cocaine on dopamine release, 50-kHz USV, and locomotor activity in rats suggests that it may also be effective to attenuate the reinforcing, cue-evoked craving, and psychostimulant effects of cocaine in humans.

According the 2020 UNODOC World Drug Report informs 65 million people in the world consume stimulant drugs. Therefore, it is crucial to find putative drugs to treat stimulant use. The results of the present study suggest that GABA_A_-R positive modulators such as benzodiazepines are promising drugs to attain this need.

## Abbreviations

NAc: nucleus accumbens
VTA: ventral tegmental area
FSCV: fast scan cyclic voltammetry
USV: ultrasonic vocalizations

## ACKNOWLEDGEMENTS

We thank Delegado Víctor A. Lopez from the Superintendência da Polícia Federal of Paranaguá for the donation of cocaine and Dr. Guilherme L. Sassaki for determining cocaine purity. We also thank Dr. Alexander Gómez-A for assistance with statistical analysis of the electrochemical data and Dr. Charles Blaha for the use the MINCS and WINCS instrumentation for FSCV experiments. This study was financed in part by the Coordination for the Improvement of Higher Education Personnel – Brazil (CAPES, PROBRAL GRANT 88881.198683/2018-01) and by The Brazilian National Council for Scientific and Technological Development (CNPq, grants 432061/2018-5, 306855/2017-8, 465346/2014-60, and: 312201/2019-2), The Brazilian Financier of Studies and Projects (FINEP, grant 01.19.0153-00), Deutsche Forschungsgemeinschaft (DFG, SCHW 559/15-1); Deutscher Akademischer Austauschdienst (DAAD, 57446778).

## AUTHOR CONTRIBUTIONS

CDC conceived the project, acquired funding, planned the FSCV experimental design with DLR and WNS and the USV experimental design with RA and RKS. WSL performed the FSCV experiments and analysis with CDC and DLR. WSL also performed the USV experiments and analysis with ME. The histology and electrode preparation were done by LFJ. The analysis of locomotion was performed by JAP. All authors discussed the results. The manuscript was written by WNS and CDC. All authors commented on the manuscript.

## DECLARATION OF TRANSPARENCY AND SCIENTIFIC RIGOUR

This Declaration acknowledges that this paper adheres to the principles for transparent reporting and scientific rigour of preclinical research as stated in the BJP guidelines for Design & Analysis and Animal Experimentation, and as recommended by funding agencies, publishers, and other organizations engaged with supporting research.

